# TEAL-seq: Targeted Expression Analysis Sequencing

**DOI:** 10.1101/2024.11.26.625462

**Authors:** Georgia Doing, Priya Shanbhag, Isaac Bell, Sara Cassidy, Efthymios Motakis, Elizabeth Aiken, Julia Oh, Mark D. Adams

## Abstract

Metagenome sequencing enables genetic characterization of complex microbial communities. However, determining the *activity* of isolates within a community presents several challenges including the wide range of organismal and gene expression abundances, the presence of host RNA, and low microbial biomass at many sites. To address these limitations, we developed “targeted expression analysis sequencing” or TEAL-seq, enabling sensitive species-specific analyses of gene expression using highly multiplexed custom probe pools. For proof-of-concept we targeted about 1,700 core and accessory genes of *Staphylococcus aureus* and *S. epidermidis*, two key species of the skin microbiome. Two targeting methods were applied to laboratory cultures and human nasal swab specimens. Both methods showed a high degree of specificity, with >90% reads on target, even in the presence of complex microbial or human background DNA/RNA. Targeting using molecular inversion probes demonstrated excellent correlation in inferred expression levels with bulk RNA-seq. Further, we show that a linear pre-amplification step to increase the amount of nucleic acids for analysis yielded consistent and predictable results when applied to complex samples and enabled profiling of expression from as little as 1 ng of total RNA. TEAL-seq is much less expensive than bulk metatranscriptomic profiling, enables detection across a greater dynamic range, and uses a strategy that is readily configurable for determining the transcriptional status of organisms in any microbial community.

**IMPORTANCE:** The gene expression patterns of bacteria in microbial communities reflect their activity and interactions with other community members. Measuring gene expression in complex microbiome contexts is challenging, however, due to the large dynamic range of microbial abundances and transcript levels. Here we describe an approach to assessing gene expression for specific species of interest using highly multiplexed pools of targeting probes. We show that an isothermal amplification step can allow profiling of low biomass samples. TEAL-seq should be widely adaptable to the study of microbial activity in natural environments.

## INTRODUCTION

As microbiome research matures, studies are shifting from those that establish correlations between microbial profiles and disease states to those that define the mechanisms by which microbes impact host physiology (1, 2). Metagenome shotgun sequencing (mWGS) analysis has been a powerful discovery tool, but it cannot address the functional state of constituent microbes. Measurement of microbial gene expression in a natural context is key to understanding the mechanistic forces driving commensalism and pathogenesis within community and host relationships (3–6).

Changes in diet, drug treatment, introduction of a pathogen, or alteration in host pathways may change the activity of the microbiome, not just its composition (6, 7). Metatranscriptome (metaTx) profiling can uncover which organisms are transcriptionally active, and which genes are expressed. However, the efficacy of metaTx is limited by the large dynamic range of organismal abundance, gene expression levels, and—in some cases—the presence of host transcripts. Thus, quantitation of gene expression in lower abundance organisms by untargeted metaTx sequencing remains infeasible due to the cost of sequencing to sufficient coverage depth. A few metaTx studies have been published that illustrate this point. One study evaluated the contribution of vitamin B12 production by skin microbes to the development of acne (8). Another examined transcription of the fecal microbiome of adult men (9). Both studies required ∼100 million reads/sample. New, more accessible methods are needed to transcriptionally profile microbes of interest in their natural milieu.

The skin is an ideal proving ground for the development of new technologies that measure microbial gene expression in mixed communities. The abundance of bacterial species can vary widely at different skin sites but importantly, abundance is a poor prognosticator of functional importance (10–12). At most skin sites host cells contribute over 90% of the total extracted nucleic acids, making mWGS and metaTx extremely inefficient, particularly for characterizing lower abundance organisms (13). *Staphylococcus (S.)* spp. are keystone contributors to cutaneous immunity, barrier integrity, and microbial community homeostasis, including antagonism with skin pathogen *S. aureus* (14–17). Our previous metaTx work with skin swabs, however, yielded <5% of reads corresponding to microbial transcripts, highlighting the challenges of studying these communities using common methods (unpublished).

Targeting microbial profiling to specific genes or genomes enables more cost-effective and comprehensive evaluation of microbial composition and activity. Array-based (18–20) and capture-based (21–24) strategies have been used to assess bacterial and viral abundance, but those methods are either difficult to customize, expensive, laborious, or all three. The availability of low-cost oligonucleotide pools from multiple vendors offers assay development options based on highly multiplexed custom-designed probes, including the ability to iteratively optimize pools based on performance and facile incorporation of new content.

We previously developed MA-GenTA (Microbial Abundances from Genome Tagged Analysis) as a quantitative and cost-effective method for species-level microbial profiling (25). We used highly multiplexed (>16,000) oligonucleotide probes to derive relative abundance data for microbes from the mouse gut. MA-GenTA and mWGS data demonstrated excellent correlation down to 0.01% relative abundance (25) and enabled inference of gene pathway composition at a cost only modestly higher than 16S rRNA amplicon sequencing. Here we build on this experience by extending the methods to targeted gene expression profiling.

## RESULTS

### Experimental design

We selected *Staphylococcus aureus* (*Sa*) and *S. epidermidis* (*Se*) for developing targeted transcriptome profiling methods as they are keystone species in the skin microbiome, important for human health and disease, and well-characterized genomically and functionally (26–28). Two targeting strategies were pursued: a single-primer extension (SPE) method based on MA-GenTA (25) and a molecular inversion probe (MIP) method, previously published for high-plex SNP genotyping assays (29). SPE uses a single 40-base primer that anneals to target sequences to prime DNA synthesis on a partially constructed Illumina sequencing library. MIP involves hybridization of the probe to the target, followed by extension across the gap region, ligation, and amplification of closed circles using primers with Illumina-compatible tails. Both methods target a cDNA or genomic DNA template.

We established experimental conditions to evaluate each targeted sequencing approach. Initial tests used RNA and genomic DNA isolated from pure cultures of *Se, Sa*, mixtures of the two, and controls that should not contain target sequences. We compared technical reproducibility, sensitivity to RNA input amount, specificity of targeting in mixed samples, the impact of rRNA depletion, the effect of pre-amplification of cDNA prior to capture, and comparison of targeted RNA-seq methods with standard bulk RNA-seq (**Figure 1**).

**Figure 1.**
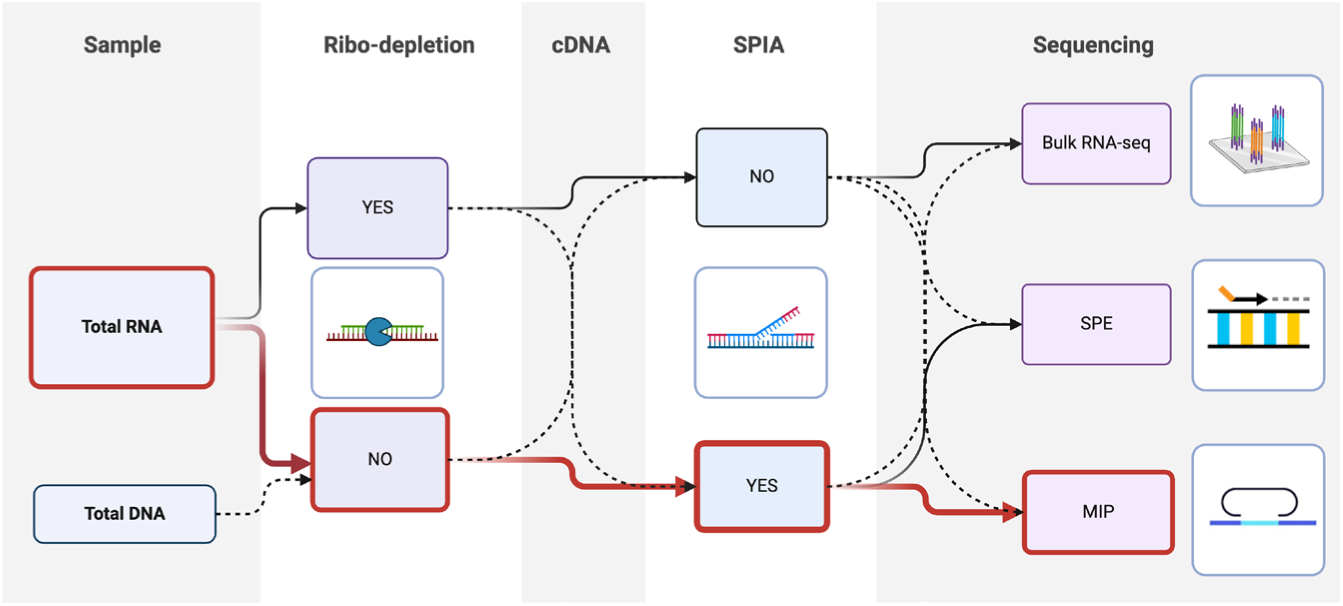
Layout of the experimental design. The experimental variables used for evaluating the targeted RNA-seq approaches are shown, illustrating how RNA samples were processed for analysis to determine the optimal TEAL-seq method.

### Probe pools are >94% on-target and species-specific

Probe design for each species maximized within-species binding and limited cross-species binding (see Methods). The SPE design encompassed 6,121 probes targeting 1,723 *Sa* coding sequences (CDS) and 5,879 probes targeting 1,670 *Se* CDS (**Figure 2A**). The MIP design encompassed 4,960 probes targeting 1,694 *Sa* CDS and 4,863 probes targeting 1,748 *Se* CDS (**Figure 2B**). A combined pool of *Se* and *Sa* probes was synthesized for each targeting method.

**Figure 2.**
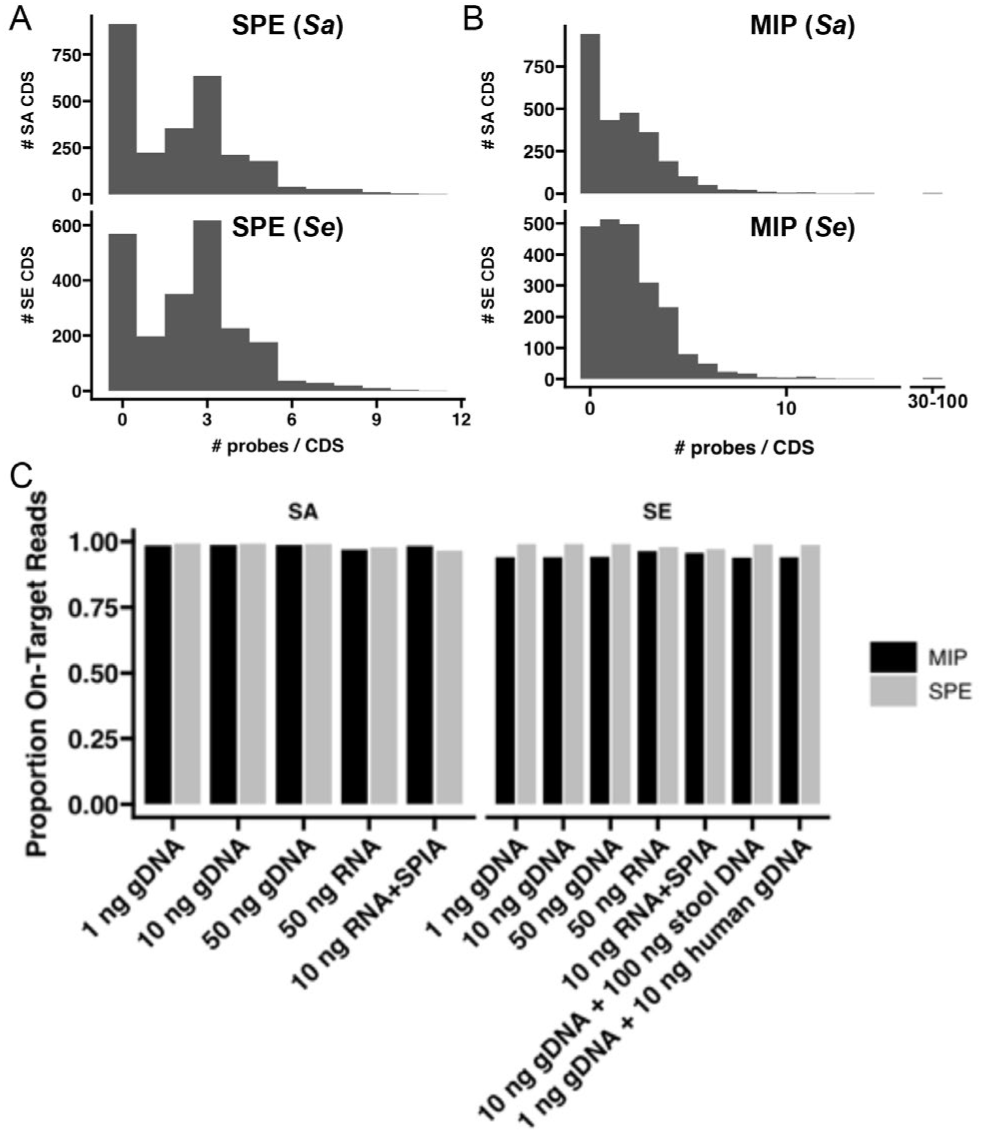
Distribution of number of probes per gene in the targeted strains. A) Probes target >50% *Sa* and *Se* CDS in SPE with at least one unique probe. B) Probes target >50% *Sa* and *Se* CDS in MIP with at least one unique probe. C) A high proportion of reads map on target with when tested on DNA and RNA controls.

Given the low biomass of the microbiome at many body sites, we evaluated the use of single-primer isothermal amplification (SPIA) (30) to increase the amount of template available for targeting and library construction. A theoretical advantage of targeted sequencing is that it makes depletion of rRNA unnecessary; we therefore also tested the effect of rRNA depletion on both bulk and targeted RNA-seq.

For both SPE and MIP, 94-99% of reads map to the targeted regions across all experimental conditions, indicating a high efficiency of targeting (**Figure 2C**, Supplemental Table 1). Notably, no ribosomal (r)RNA reads were detected, even in samples processed without rRNA depletion. Single-species experiments showed that probes were highly specific to the targeted genome when full length reads were aligned to the reference genome. In addition, few reads were obtained from *E. coli* RNA, human genomic DNA, and mouse microbiome RNA (cDNA) (data not shown), indicating that cross-hybridization is minimal, even in the presence of complex microbial nucleic acids.

### Targeted transcriptomics is reproducible

We tested the reproducibility of inferred gene expression levels based on counts per million (CPM) per probe using MIP and SPE by comparing variation across technical replicates to that of bulk RNA-seq. We found technical replicates (libraries prepared from the same RNA sample) highly correlated for all sequencing approaches (**Figure 3A**). Subsequently, we determined that libraries from the same RNA sample with and without SPIA treatment were highly correlated for bulk RNA-seq and MIP, but less so for SPE (**Figure 3B**). Further, when SPIA was used, libraries prepared from standard amounts of RNA (“Standard Input”, 10 ng) and an order of magnitude less RNA (“Low Input”, 1 ng) were still correlated for both MIP and SPE (**Figure 3C**).

**Figure 3.**
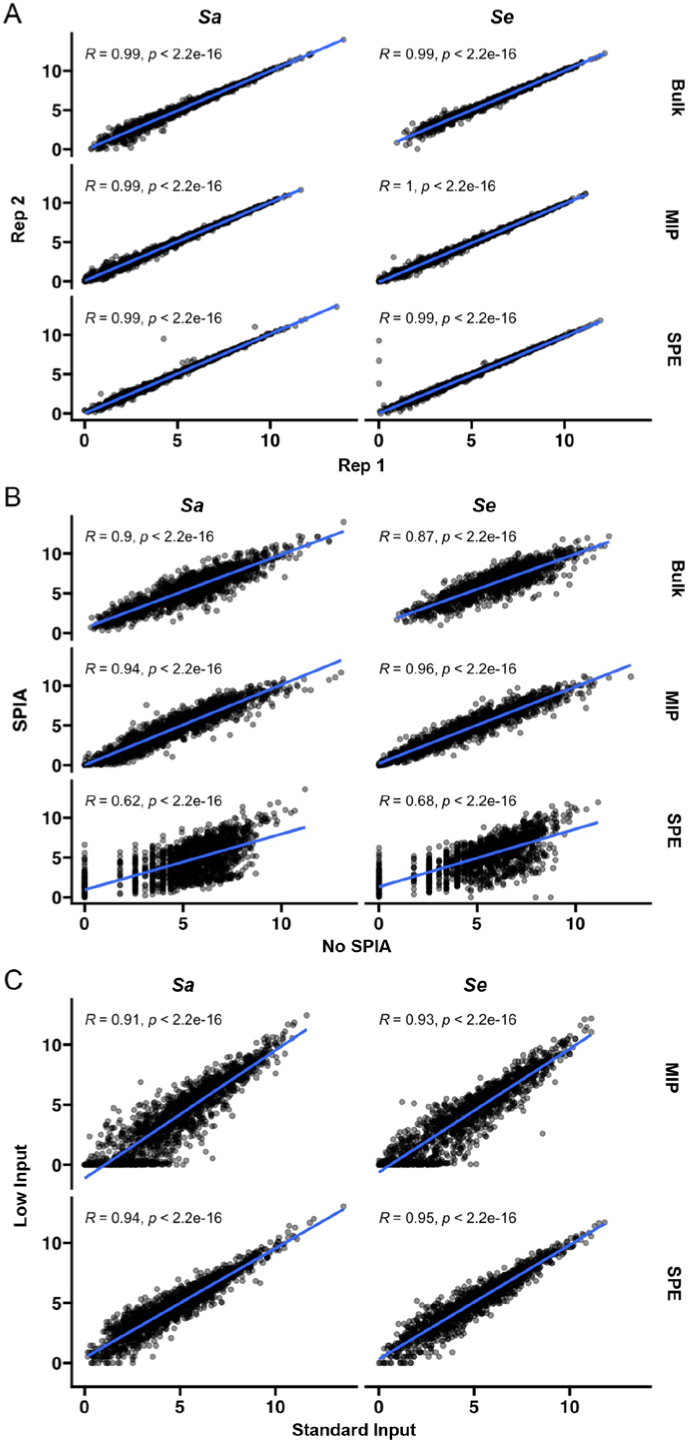
Evaluation of reproducibility using a 50:50 mixture of *Sa* and *Se* RNA. A) Technical replicates (n=2) are highly correlated for bulk RNA-seq, MIP and SPE. B) Samples with and without SPIA have good correlation for bulk RNA-seq and MIP, but less so for SPE. C) Samples with low (1 ng) and standard (10 ng) input total RNA amounts correlate well in MIP and SPE. Correlation coefficients (*R*) and p-values from Pearson tests.

### Probe performance varies across CDS

Targeted sequencing of *Se* and *Sa* gDNA was used to evaluate the extent of variation in probe performance. We observed a wide range of CPM/probe, with *Se* probes having a much broader range than *Sa* probes in both the SPE and MIP datasets (**Figure 4A**). Since *Se* and *Sa* probes were synthesized as a single pool, there is no *a priori* reason why they should have such different performance between the two species. We considered whether there is a gradient of genomic copy number ranging from the origin to the terminus of replication due to differences in growth rate (31), however, we found that genome position was poorly correlated with CPM levels in both species (Supplemental Figure 1).

**Supplementary Figure 1.**
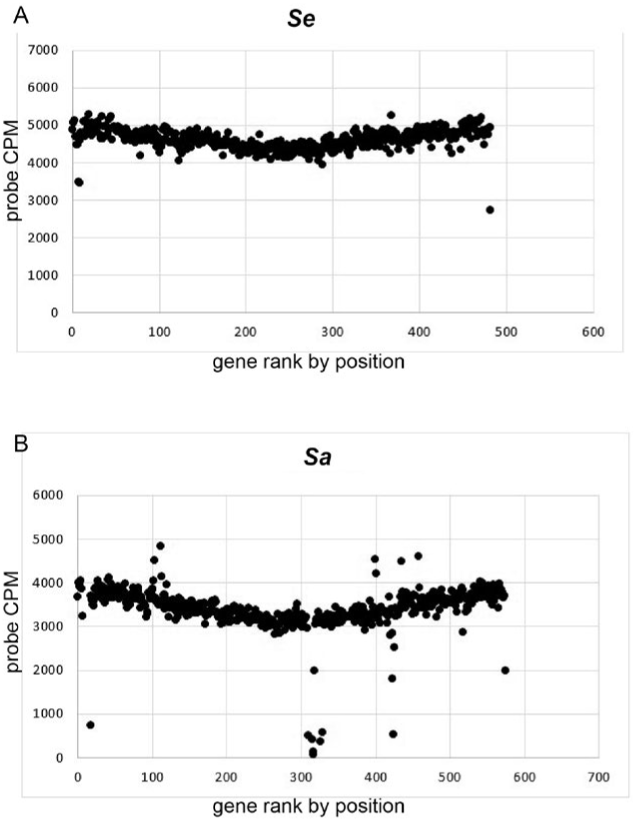
CPM bias based on gene position relative to origin of replication. There is no substantial bias introduced by position relative to the origin of replication for *Se* Tu3298 (A) or *Sa* USA300 (B).

**Figure 4.**
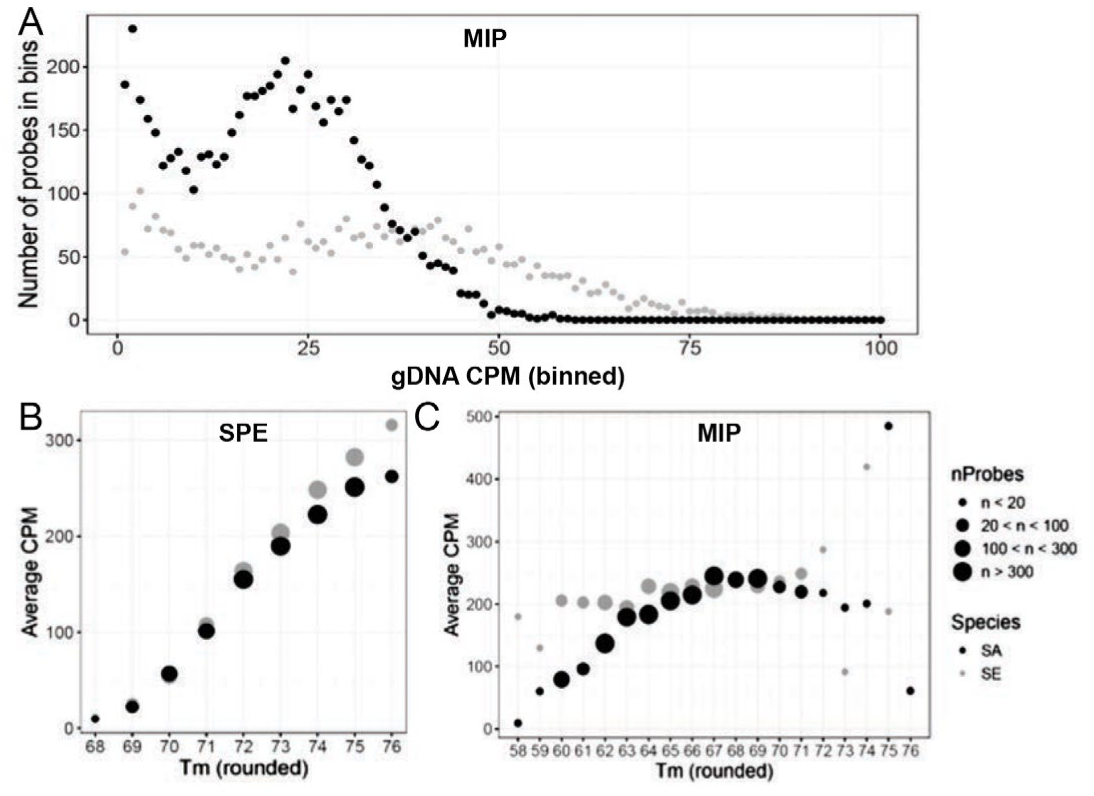
Probe performance varies and correlates with melting temperature in SPE data more so than MIP data. A) Sequencing of gDNA with SPE shows a range of CPMs for *Sa* (black) and *Se* (grey) probes. B) Probe CPM correlates with melting temperature in SPE data for both *Sa* (black) and *Se* (grey). C) CPM has minimal correlation with melting temperate in MIP data for both *Sa* (black) and *Se* (grey).

For SPE probes, there was a marked increase in average CPM with probe melting temperature (T_M_) (**Figure 4B**), whereas T_M_ had minimal correlation with CPM for MIP probes (**Figure 4C**). We also considered the performance of probes that have mismatches to the test genome compared to perfect-match probes. This is an important consideration for profiling of microbial communities, because the exact sequence of the target genomes will not often be known. SPE probes showed a substantial reduction in read counts with increasing numbers of mismatches, whereas MIP showed little or no effect of up to three mismatches (Supplemental Figure 2). Given the limitations of the SPE approach, we focused our remaining analyses on the MIP datasets.

**Supplemental Figure 2.**
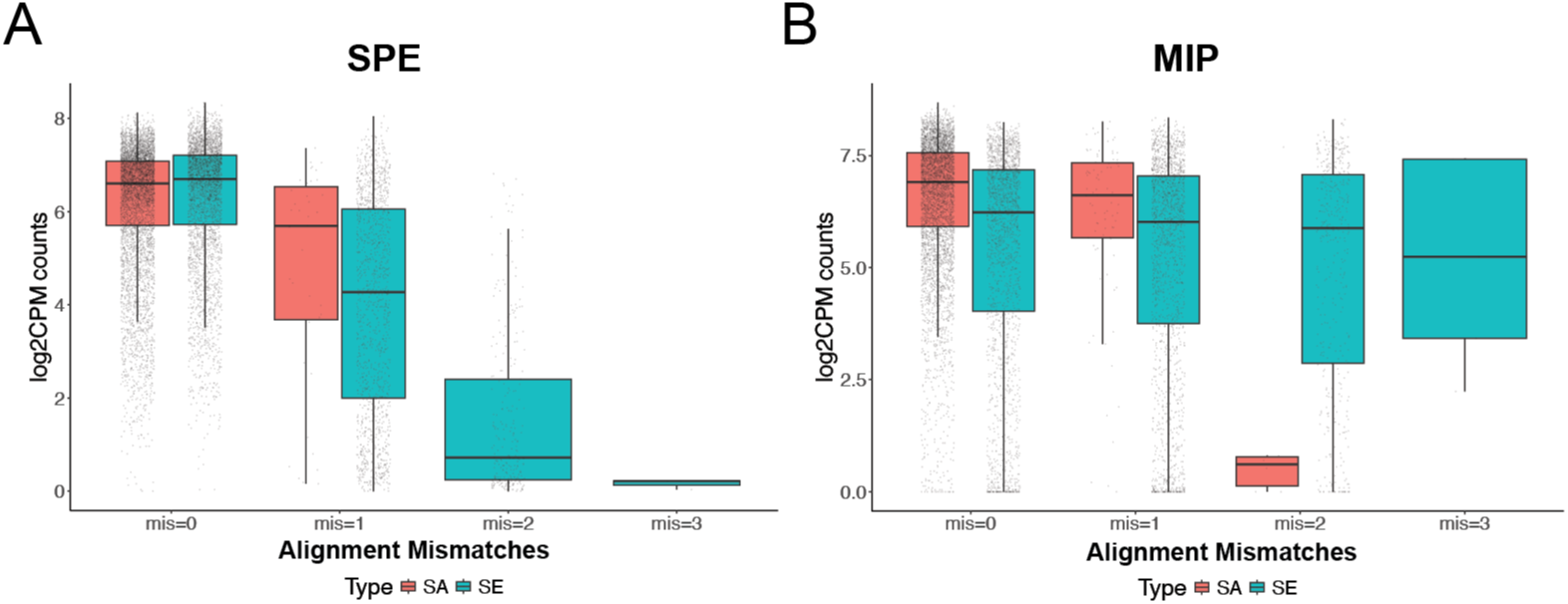
Read mapping to experimental genomes. Mapping rates as the number of mismatches between the probe design sequence and the experimental genome increase for SPE (A) and MIP (B).

### Targeting strategies are compatible with signal amplification

The inability to perform oligo-dT priming for cDNA synthesis from microbial transcripts means that rRNA must be depleted as part of library construction for bulk RNA-seq. However, targeting offers the possibility of using total RNA as input, thus reducing processing steps and any potential bias introduced by rRNA depletion. We tested the fidelity of targeting (MIP) with and without ribodepletion and compared the results to bulk RNA-seq. There is only slightly lower correlation in CPM between samples sequenced by bulk RNA-seq and MIP (mean R^2^=0.47, **Figure 5A** for *Sa*, **Figure 5B** *Se*) than there are between MIP samples with and without ribo-depletion (mean R^2^ = 0.64, **Figure 5C** for *Sa*, **Figure 5D** *Se*), indicating that rRNA depletion had a substantial impact on many non-ribosomal transcripts.

**Figure 5.**
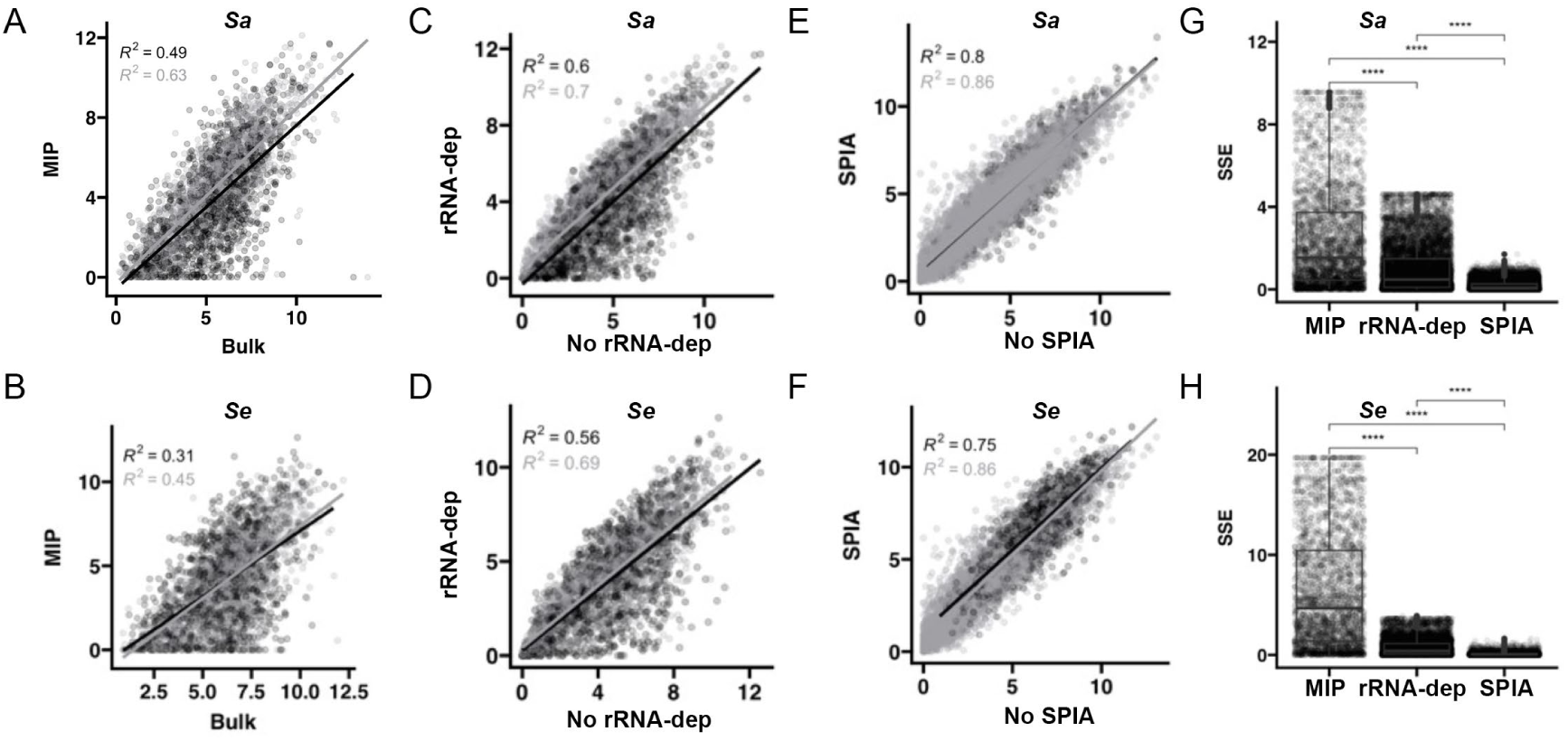
Sources of variation in targeted RNA-seq. A) comparison of CDS-level expression in bulk RNA-seq and MIP (targeted) without SPIA (black) and with SPIA (grey) for *Sa* and *Se* (B). C) MIP with and without RNA depletion without SPIA (black) and with SPIA (grey) for *Sa* and *Se* (D). E) MIP with and without SPIA amplification with ribo-depletion (black) and with ribo-depletion(grey) for *Sa* and *Se* (F). G) Quantification of sum of squared error (SSE) across comparisons for *Sa* and *Se* (H). Correlation coefficients (*R*) and p values from Pearson tests. **** p<0.0001, Wilcoxon test.

To assess SPIA’s influence on resulting mRNA measurements, we quantified mRNA abundance as inferred from bulk RNA-seq versus MIP with and without SPIA. Bulk RNA-seq and MIP samples with and without SPIA correlated highly (mean R^2^ = 0.83, **Figure 5E** for *Sa*, **Figure 5F** *Se*), more highly than samples with and without rRNA depletion. Further, CDS-level CPM values across bulk and targeted RNA-seq were less variable when SPIA amplification was used (**Figure 5A**, **B** grey) than when SPIA was not used (**Figure 5A**, **B** black). In summary, since SPIA amplification introduces significantly less sum squared error (SSE) than ribo-depletion does (p<0.0001 by Wilcoxen test) for both species (**Figure 5G and 5H**), we determined MIP with SPIA to be the optimal workflow for TEAL-seq (as outlined in **Figure 1**).

### TEAL-seq captures gene expression changes across growth conditions

A major goal of RNA-seq analysis is to identify differentially expressed genes (DEGs) across conditions of interest to infer underlying biology and nominate targets for downstream functional studies. To test the ability of TEAL-seq to capture DEGs under physiologically relevant conditions, we measured gene expression from *Sa* and *Se* cultures grown in TSB (pH 7), acidic TSB (pH 4.8), and TSB with urea (4.5%w/v) to model acid stress commonly encountered by staphylococci *in vivo*. Growth conditions were delineated by the first and second PCs which together explained 97% of variance for *Sa* (**Figure 6A**) and 94% of variance for *Se* (**Figure 6B**). Further, genes with the highest fold-change in response to acid were consistent with known acid responses, in particular urease genes in *Sa* (**Figure 6C**) and genes previously identified as upregulated by acid in *Se* (**Figure 6D**). Clustering of *Sa* samples shows that the expression patterns are driven first by condition (**Figure 6E**). For *Se*, condition also influences expression of these genes of interest, although urea treatment had less of an effect than acid (**Figure 6F**). Our results suggest that TEAL-seq performs predictably for the analysis of differential gene expression and thus is a valuable approach for comparative microbial studies.

**Figure 6.**
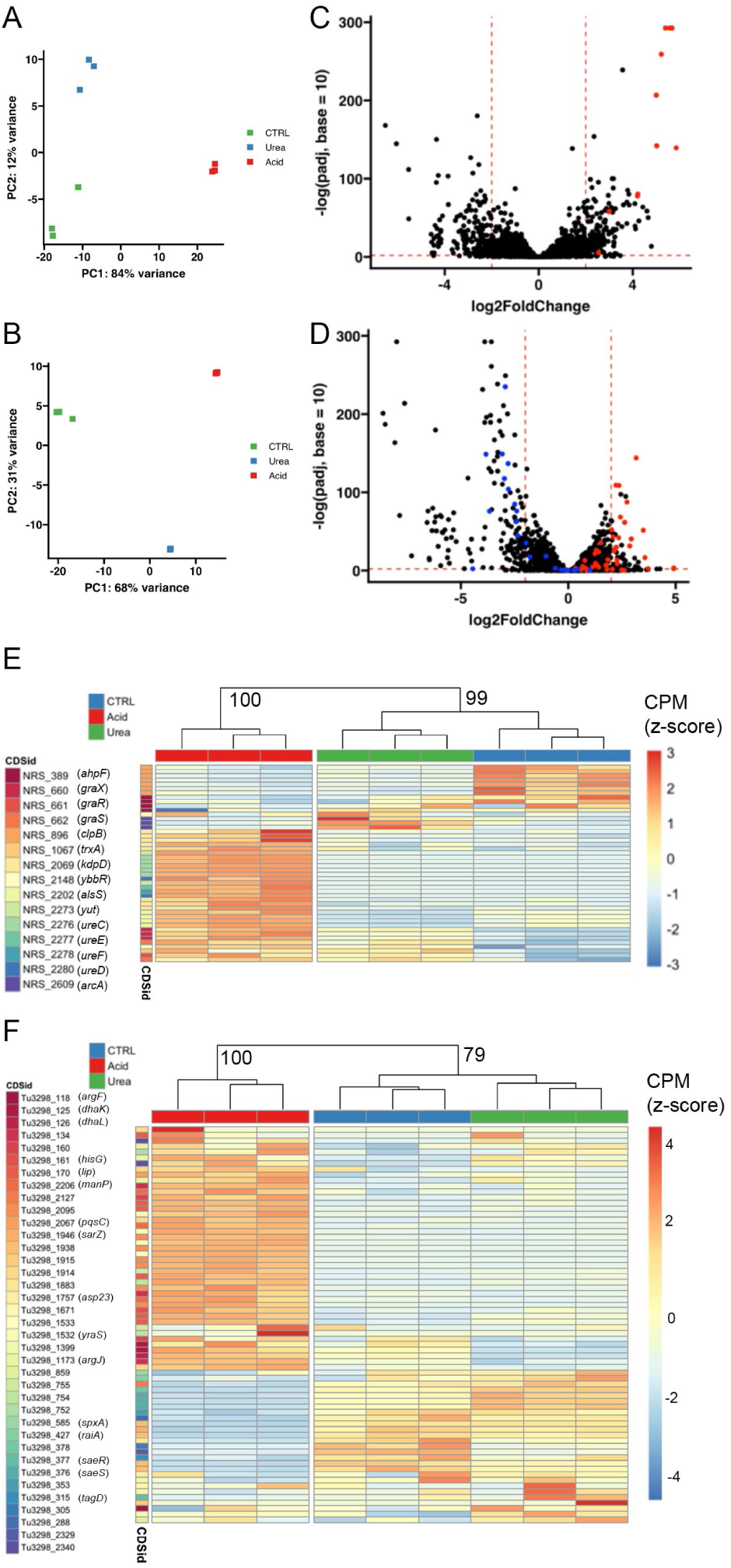
TEAL-seq detects urease and acid response genes as differentially expressed under acid stress. A) Principal components of targeted gene expression separate samples by growth condition for *Sa* and *Se* (B). C) Differential expression between control (TSB) and acid treatment identifies urease genes for *Sa (*red) and the previously published acid response for *Se* (red and blue) (D). E) Urease and acid response probes across conditions are consistent across CDSs for *Sa* and *Se* (F). Each column represents biological replicates (n=3).

### DEG analysis from model and human samples

To model the technical challenge of low amounts of material extractable from skin microbiomes, we applied TEAL-seq to RNA extracted from reconstructed human epidermis (RHE) colonized with clinical skin microbiome isolates as well as RNA extracted directly from human nasal swabs. *Sa* and *Se* transcripts were readily detected from both samples. Transcript levels derived from these sources were substantially different from those grown in TSB (PC1, separation by sample type accounted for 37% of variance for *Sa* (**Figure 7A**) and 46% of variance for *Se* (**Figure 7B**). Notably, samples from independent RHE and swabs were more variable than replicates of growth in TSB, as expected for complex models and *ex vivo* samples. Further, there was high correlation in expression profiles for samples grown in TSB or between sequencing replicates of the same sample for both *Sa* (**Figure 7C**) and *Se* (**Figure 7D**). Thus, this variation is likely biological.

**Figure 7.**
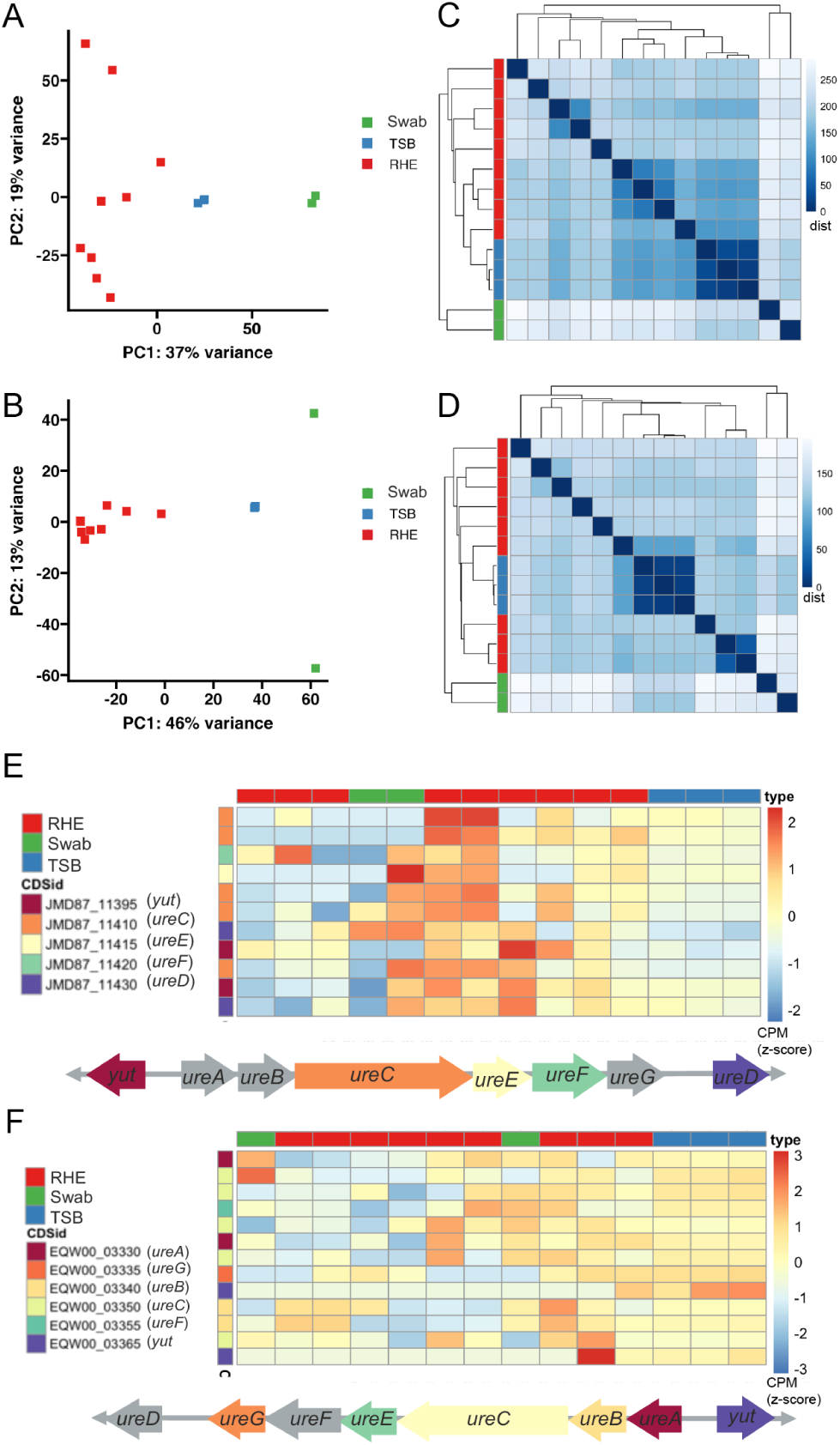
*Sa* upregulates urease gene expression when grown on RHE and as sampled in nasal swabs, but *Se* does not. A) PCAs separate expression from growth on RHE from *in vitro* (TSB) and human nasal swabs in *Sa* B) and *Se.* C) Samples correlate by source more than by SPIA treatment in *Sa* D) and *Se*. Values are pairwise distances in Pearson correlation matrices. E) Urease genes (operon structure and direction per diagram) are induced on RHE and in nasal swabs for *Sa* F) but not *Se*.

We used TEAL-seq to compare expression patterns of urease genes from bacteria grown in TSB, on RHE, or isolated from swab samples. A subset of both nasal swabs and RHE samples showed marked increase in the expression of urease genes relative to TSB for *Sa* (**Figure 7E**). Interestingly, another subset of swab and RHE samples had lower expression of urease genes compared to TSB for *Sa*. *Se* samples from RHE and swabs also displayed variable urease gene expression compared to those grown in TSB, although the trend was less pronounced than in *Sa* samples (**Figure 7F**). To further validate TEAL-seq as a faithful measure of mRNA in complex samples, we attempted to compare the expression of urease genes by TEAL-seq to that detectable by bulk RNA sequencing on the same RHE sample. Notably, not all samples had detectable microbial mRNA using bulk RNA sequencing; colonized RHE samples sequenced to depths of 8.5 – 42.1 million (m=30.4 million) contained only 0.06 – 1.5% (m=0.3%) reads that mapped to *Sa* or *Se* mRNA and thus failed to provide sufficient reads to measure genome-wide expression (Supplemental Table 3). These results demonstrate that TEAL-seq can be used to measure gene expression of *in vivo* relevant pathways in complex samples in a manner more sensitive than the current state-of-the-art method.

## Discussion

We present an approach for targeted gene expression analysis optimized for samples with low biomass—TEAL-seq—that yields a high proportion of reads on target, even in the presence of total RNA input and complex background RNA or DNA. We tested two oligo-based methods for targeting, MIP and SPE, and determined MIP to be more robust than SPE. Both methods were capable of inference of mRNA levels relative to bulk RNA-seq, but expression levels inferred from MIP had less probe-to-probe variation. Further, we showed that cDNA amplification using SPIA reduced variation between replicates and had less impact on inferred expression level than rRNA depletion. Therefore, we propose the use of MIP with SPIA for measuring bacterial transcript levels in low biomass samples.

We showed that probes designed based on genes of laboratory strains captured gene expression information from other common experimental strains and clinical isolates of the same species. This is in part due to short probe length, ensuring coverage of conserved genes across strains, as well as the design, which enabled integration of data across multiple probes per CDS. Cross-strain capture is an important feature of this method as the application of TEAL-seq to measure gene expression of real-world samples will require probes to be designed from existing genomic or metagenomic data. Our benchmarking of results from probes designed for *Sa* strain M2872 and *Se* strain ATCC14990 and applied to *Sa* strain USA300 and *Se* strain Tu3298 as well as clinical isolates of both species demonstrate the flexibility of TEAL-seq.

We used TEAL-seq to assess changes in gene expression of *Sa* and *Se* grown in TSB at pH 7 and pH 4.8 and identified highly concordant changes in expression compared to previous studies. We confirmed that *Sa* responds to acid stress with urease genes and *Se* does not. This is significant as urea is a human skin-produced metabolite that has important roles in barrier defense (32), and differences in *Sa* and *Se* responsiveness to acidic conditions may explain differences in invasiveness (33).

MetaTx has had limited usage in studying low biomass microbiomes. However, TEAL-seq successfully captured predictable expression profiles from multiple complex sample types, including those isolated clinically. New tools are needed for identification of population-level expression of virulence pathways or immunogenic metabolites at species resolution, as these will contextualize the function of the microbiome as a collective and refine our understanding of the contribution of individual species to the community (34–37). Additional studies applying TEAL-seq in the context of complex natural microbial communities will be needed to further demonstrate the potential of this method in experimentally relevant systems. However, the robust performance of TEAL-seq in capturing biologically significant transcriptional patterns from skin microbes grown under skin-relevant stress and reconstructed human epidermis make it a promising tool for future applications.

## Methods

### Bacterial samples and RNA and DNA isolation

Single colonies of *Staphylococcus aureus* USA300 and *Staphylococcus epidermidis* Tu3298 were inoculated into 2ml Tryptic Soy broth (TSB) (BD cat:211825) and grown overnight at 37°C and 200 rpm agitation. Overnight cultures were sub-cultured into fresh 2 ml TSB medium at OD_600_ ∼0.05 and allowed to grow to OD_600_ ∼0.5-0.6. For stress conditions, as in (38), overnight cultures were grown as described above and then sub-cultured into mildly acidic TSB (pH 4.8) or TSB+ 4.5% w/v urea and allowed to grow to OD_600_ ∼0.5-0.6. For RNA extraction, to 1 ml of each mid-log phase culture, 2 ml of RNAprotect® Bacteria Reagent (cat # 76506, Qiagen Inc.) was added, and cells were centrifuged at 8000 x g for 5 minutes at 4°C. Cell pellets were then resuspended in RLT buffer (RNeasy Mini kit, (cat # 74104, Qiagen Inc.) and transferred to 2 ml safe-lock tubes containing 50 mg acid-washed glass beads (150– 600 μm diameter). Cells were homogenized using a TissueLyser for 4 minutes at 30Hz/sec and further processed according to the kit protocol. For DNA extraction, cells from 1 ml of culture were pelleted by centrifugation at 16000 x g for 2 minutes and DNA extracted using GenElute™ Bacterial Genomic DNA Kit (cat# NA2110, Sigma-Aldrich).

For mixed cultures, overnight cultures of individual strains were sub-cultured as described above and allowed to grow to an OD_600_ ∼ 0.5-0.6 in TSB at 37°C with 200 rpm agitation. Equal numbers of cells from each strain were then inoculated into fresh 5 ml TSB at an OD_600_ ∼0.1-0.2 and grown at 37°C with agitation at 200rpm for 2-3 hours until the OD_600_ reached ∼0.5.

Mouse microbiome DNA was prepared as previously described (39). RNA was extracted from bacteria grown in association with a reconstructed human epidermis (RHE) skin organoid model (MatTek EpiDerm) with the Qiagen RNeasy kit (cat # 74104, Qiagen Inc.) and *Escherichia coli* RNA was purchased from ThermoFisher (#AM7940).

### Reconstructed human epidermis cell culture cultivation

9mm primary normal human RHE tissue cultures were obtained from MatTek Corporation (EpiDerm, Gothenburg, Sweden). All batches were grown using cells from MatTek Corporation’s EpiDerm standard donor, a healthy male. The RHE were cultured according to the manufacturer’s directions. Briefly, upon arrival, the RHE cultures were placed in 6-well plates with 1mL of warmed antibiotic-free EpiDerm Maintenance Media (MatTek Corporation,) or the equivalent EpiDerm Assay Media basally per well. The basal media was replaced daily, and the RHE cultures were kept at 37°C with 5% CO_2_.

For each microbial isolate, a single colony was grown overnight in sterile 1X TSB. 10^8^ colony-forming units from each liquid culture were washed with ultrapure water (Fisher Scientific, #AAJ71786AP, Hampton, NH) and resuspended to a final concentration of 10^7^ colony-forming units in 120 µL.

The RHE cultures were then dosed with 120uL of microbial isolate or vehicle (ultrapure water alone). Dosed RHE cultures were incubated for 1 hour at 37°C, then inoculum was aspirated to restore the air liquid interface. RHE were incubated with remaining bacteria for 18 hours prior to harvest.

At harvest, 200 µL of PBS (MatTek Corporation) was added to the apical surface of each RHE culture, pipette mixed, removed, and plated for CFU enumeration. 140 µL of RLT buffer + 1% beta-mercaptoethanol was added to each RHE culture for RNA preservation. The RLT buffer-tissue solution was frozen at-80°C until RNA extraction. *S*. *epidermidis* Tü3298-GFP colonized RHE were visualized under blue light.

### Target gene selection and probe design

Pangenome datasets for *Sa* and *Se* were assembled using Roary (40) based on the available complete genome sequences of each species: 51 complete *Sa* and 86 complete *Se* genomes downloaded from GenBank. CDS present in at least 50/86 *Se* and 30/51 *Sa* genomes were submitted for probe design. Candidate probe sets were designed using standard informatics workflows at Tecan Genomics for SPE probes and Molecular Loop for MIP probes. We selected probes for inclusion in the custom pool by subsampling based on perfect BLAST matches to at least 50 *Se* or 30 *Sa* genomes and exclusion of probes with alignments having fewer than five mismatches to the alternative species or to a database of 10 *S. capitis* and 15 *S. hominis* genomes. Additional filtering criteria removed probes at the extremes of melting temperature and sequence composition (requiring no runs of >6 of a single base) and established minimum probe spacing of at least 100 bases. All probes targeting both genomes were synthesized and used in a single pool. Distributions of the number of probes per CDS is provided in Figure 2. Probe sequences are in Supplemental Table 2. Purified nucleic acids were split across several workflows as shown in Figure 1.

### Bulk RNA-seq library preparation and sequencing

The NEBNext Ultra II Directional Library Prep protocol (NEB #E7760S) was used with NEB’s bacterial rRNA depletion kit (NEB # E7850S) to make bulk RNA-seq libraries for whole transcriptome sequencing. 100 ng RNA input was used and adaptors were diluted 25-fold before ligating to the cDNA. The same set of RNA samples was used in a hybrid workflow combining pre-amplification with the NEBNext Ultra II Library Prep. 10 ng of total RNA was subjected to rRNA depletion and then cDNA synthesis and amplification using the Single Primer Isothermal Amplification (SPIA) method with the Crescendo cDNA synthesis for qPCR kit (Tecan Genomics, Inc.). For these samples, RNA was not fragmented prior to cDNA synthesis. After SPIA amplification, cDNA was fragmented using components of the Allegro Targeted Genotyping V2 kit (Tecan Genomics, Inc.) before proceeding with end repair, adaptor ligation (25-fold diluted adaptors) and PCR-enrichment using the NEBNext Ultra II Directional Library Prep kit components. *E. coli* RNA was included as a positive control in both the workflows. Index PCR was performed using the NEBNext Multiplex Oligos for Illumina (Dual Index Primers Set 1) (NEB #E7600S). The final library purification was carried out using 0.78x Ampure XP beads to remove the adaptor peak. Libraries were assessed on the TapeStation and Qubit dsDNA high sensitivity assay before normalizing to 4 nM.

### cDNA synthesis and SPIA amplification

First and second strand cDNA synthesis were performed using the Crescendo cDNA synthesis for qPCR protocol (Tecan Genomics, Inc.). cDNA samples were then divided, with one portion subjected to SPIA amplification per the Crescendo protocol. cDNA was purified with AMPure XP beads at 1.8x and eluted in 33 ul of 1x TE. The resulting cDNA samples were quantified using NanoDrop following the kit protocol and Qubit dsDNA high sensitivity assays. The samples that were amplified with SPIA yielded >1 ug of cDNA and the non-SPIA samples were 6-8 ng.

### Targeted RNA-seq library preparation and sequencing using single-primer extension probes

SPIA-amplified cDNA samples were diluted 20-fold and 10-20 ng of cDNA was used as input for the Allegro Targeted Genotyping V2 protocol (Tecan Genomics, Inc.) with the custom probe pool. Non-amplified cDNA was not diluted and ∼1 ng was used as input for the Allegro protocol. Genomic DNA was used at 10-50 ng input. Each sample was enzymatically fragmented, followed by ligation of barcoded adaptors. Barcoded samples were then purified, pooled, and placed in an overnight hybridization reaction mixture with the probe pool. The following day (>12h) the DNA polymerase enzyme was added to the reaction for extension at 72°C for 10 min. Post-enrichment purification was done with 0.8x AMPure XP beads. A qPCR step was used to determine the number of cycles for library amplification (11 cycles). The final library pool was bead purified with 0.8x AMPure XP beads. A low template control (LTC) with 50 pg of *S. epidermidis* RNA input and a mouse stool metagenome DNA sample were used as sensitivity and specificity controls respectively. The library pool was adjusted to 32 nM and 20 ul of the pool was used for sequencing (20 fmole/sample). The final sequencing pool also comprised libraries created using the NEBNext Ultra II Library prep protocol and Illumina DNA Prep (M) kit that were normalized to 4 nM and 0.2 nM respectively to represent 20 fmoles for the NEBNext samples and 1 fmole for the Illumina DNA Prep libraries. The exception was the *E. coli* RNA positive control (100 ng input) that was pooled at a lower amount to represent 5 fmoles in the sequencing pool. The library pool was loaded on NovaSeq 6000 SP flowcell 2×100bp run. The custom read 1 (R1) primer provided in the Allegro kit was spiked into the Illumina read 1 primer position at 0.3 uM and a phiX174 control library was spiked-in at 1% as recommended.

### Targeted RNA-seq library preparation and sequencing using molecular inversion probes

Targeted RNA libraries were prepared with the Low Input DNA Target Capture kit (Molecular Loop Biosciences, Inc.) using the cDNA generated with the Crescendo kit (Tecan Genomics, Inc.), including both the SPIA and non-SPIA pre-amplified samples, as well as the gDNA samples. The sample input was not normalized (50 pg – 50 ng) to evaluate its effect on probe hybridization efficiencies and consequently the number of sequencing reads generated from each of these samples. Among the 32 samples, only the human blood high molecular weight genomic DNA sample was pre-treated by heating it to 95°C for 5 min for denaturation. The protocol was followed using the kit user guide and the custom probe pool was hybridized for 18 hours (recommended ≥16 hours). After hybridization, each library was amplified by PCR for 20 cycles, pooled at equal volume (10 ul each), and the library pool was purified with 0.8x Agencourt RNAXP Clean beads and eluted in 25ul of 1xTE. The library pool was quantified with the Qubit high sensitivity dsDNA assay and on the TapeStation D1000 high-sensitivity ScreenTape, which resulted in a single peak at ∼415 bp. The library pool was loaded on a NovaSeq 6000 SP flowcell 2×100bp run. The custom workflow was selected for the sequencing run using custom read1, read2 and sequencing primers provided by Molecular Loop. The phiX174 control library was spiked-in at 1%.

### Illumina DNA Prep Libraries

Libraries of *Se* and *Sa* gDNA were prepared using the Illumina DNA Prep, Tagmentation kit (Illumina, 20018705) and indexed using IDT for Illumina DNA/RNA UD Indexes (Illumina, 20042666). Reads were assembled using CLC Genomics Workbench and the resulting assembly was compared to the reference sequences for the *Se* and *Sa* genomes, confirming that the cultured cells represented the expected genomes.

### Genome sequence alignment

Reads were processed with trimmomatic 0.39 (41) to remove low-quality reads. For bulk RNA and SPE reads, the adapters were removed from the fastq reads using Cutadapt (42) version 4.4s, using the bait sequence ‘AGATCGGAAGAG’. For MIP reads, seqtk v1.4-r122 was used (https://github.com/lh3/seqtk) to remove the five bases from the 5′-end of the sequences as *seqtk trimfq-b 5 ˂fastq˃*. Reads were aligned to reference genomes *Sa* m2872 (GCF_017329165.1) and *Se* ATCC 14990 (GCF_006094375.1) as well as *Sa* USA300 (GCF_002993865.1) *Se* Tu3298 (https://github.com/ohlab/S.epi-CRISPRi-and-RNA-Seq) using BWA-MEM (43) version 0.7.17-r1188 as *bwa mem-t 8-k 19-w 100.* The mapped reads of each of the *Se* and *Sa* species were extracted with bamtools v2.5.2 (44) with parameter *-isMapped ‘true’* and subsequently filtered with MAPQ >30 and edit distance NM <4. High-quality sequences were sorted and converted into.bam and.bed files for downstream processing by SAMtools (45) v0.7.17-r1188. For bulk RNA-seq, the read count aligned to each CDS region was computed using featureCounts v1.6.4(46) (from Subread1.6.4) with-t CDS or-t rRNA, as appropriate, and summarized as counts per CDS per million reads per kilobase CDS length (RPKM).

### Probe sequence alignment

We also used BWA-MEM to align the SPE and MIP reads to a database of the probe sequences. For MIP, we used the seqinr R package (47) to extract the binding locations consisting of the first and the last 40bps of the probe sequences and generated new fasta files with the ‘first 40bps’ and ‘last 40bps’ probe data. We aligned the reads of each sample separately to the probe sequences of each species. The mapped reads were extracted as before with bamtools v2.5.2 and subsequently filtered with MAPQ >30 and NM <1 (perfect match) in SPE and MAPQ >30 and NM <3 in MIP. The high-quality sequences were sorted and converted into bam and bed files for downstream processing by SAMtools v0.7.17-r1188. The raw counts were generated with *bedtools coverage* for each sample from the probe sequences (.bed) and cleaned read (.bam) files by bedtools v2.31.0 (48). The raw counts were CPM normalized as 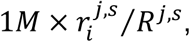 where 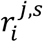 denotes the number of reads mapped to location *i* in sample *j* and species ***S***, *S* ∈ {*S*e, *S*a} and *R*^*j,s*^ is the total number of mapped reads in sample *j* and species *s*. Probe count tables are provided as **Supplemental Tables 6-10**.

### Differential Expression Analysis and Correlation Analyses

Differential expression analysis across stress conditions was performed in Rv4.4.1 (49) using DESeq2 (50) using a two factor design matrix accounting for stress treatment (TSB, acid, or urea) and SPIA status including interaction terms. In all comparisons, differentially expressed genes were defined as those with fold-change magnitude > 4 and adjusted p-value < 0.01. Correlation matrices were calculated in R as Euclidean distances using the stats package (49) and pairwise correlations were assessed via Pearson correlation coefficients (R values) of best fit linear models to two-dimensional data using ggpubr (51).

## Data availability

All data used in these analyses are available at the NCBI Gene Expression Omnibus under accession GSE279187.

## Acknowledgements

We thank the Genome Technologies Scientific Service for performing sequence data collection and Mitch Kostich for helpful discussions regarding data analysis.

## Author Contributions

GD and MA performed data analysis and wrote the manuscript. SC wrote the manuscript. PS performed the targeted sequencing experiments. IB and EM contributed to data analysis. EA performed the RHE experiments. JO and MA conceived the study.

